# Diverse microtubule-destabilizing drugs induce equivalent molecular pathway responses in endothelial cells

**DOI:** 10.1101/2025.01.22.632572

**Authors:** Lillian J. Horin, Matthew Sonnett, Boyan Li, Timothy J. Mitchison

**Affiliations:** Department of Systems Biology, Harvard Medical School, Boston, MA 02115

## Abstract

Drugs that modulate microtubule (MT) dynamics are well-characterized at the molecular level, yet the mechanisms linking these molecular effects to their distinct clinical outcomes remain unclear. Several MT-destabilizing drugs, including vinblastine, combretastatin A4, and plinabulin, are widely used, or are under evaluation for cancer treatment. Although all three depolymerize MTs, they do so through distinct biochemical mechanisms. Furthermore, their clinical profiles and therapeutic uses differ considerably. To investigate whether differential modulation of molecular pathways might account for clinical differences, we compared gene expression and signaling pathway responses in human pulmonary microvascular endothelial cells (HPMECs), alongside the MT-stabilizing drug docetaxel and the pro-inflammatory cytokine TNF-α. RNA-sequencing and phosphoproteomics revealed that all three MT destabilizers triggered equivalent molecular responses. The substantial changes in gene expression caused by MT destabilization were completely dependent on Rho family GTPase activation. These findings suggest that the distinct clinical profiles of the destabilizing drugs depend on differences in pharmacokinetics (PK) and tissue distribution rather than molecular actions. The washout rate of the three drugs differed, which likely translates to PK differences. Our data provide insights into how MT destabilization triggers signaling changes, potentially explaining how these drugs induce cell cycle re-entry in quiescent cells and how plinabulin ameliorates chemotherapy-induced neutropenia.

**Significance Statement:** Microtubule (MT)-destabilizing drugs, despite their potential for toxicity, are used to treat a surprising range of diseases, including cancers, inflammatory disorders, and parasitic infections. To investigate how drugs with apparently similar actions on MTs achieve diverse clinical effects, we compared the molecular pathways that are activated by three drugs with different clinical profiles, vinblastine, combretastatin A4, and plinabulin. All three elicited similar gene expression responses via Rho GTPase activation. This finding suggests that their distinct clinical effects are not caused by different effects on MTs, but rather by differences in drug transport, pharmacokinetics or tubulin isotype affinity. Our findings provide insights into how plinabulin might protect the bone marrow and may help medicinal chemists design MT drugs for new applications.

## Introduction

Microtubules (MTs) and their subunit tubulin (Tb) are key drug targets in research and medicine. Tb was discovered as the target of colchicine (1), an ancient medicine still commonly prescribed to treat inflammatory diseases (2). Vinca alkaloids and taxanes are mainstays in cancer treatment, alongside two new classes, epothilones and eribulin (3, 4). Additionally, several colchicine site-binding drugs are under development as tumor vascular disruptors for use with chemotherapy (5). MT stabilizers and destabilizers have been proposed as treatments for neurological diseases and spinal cord injury, though none are yet approved (6–8).

A long-standing puzzle is why different drugs that bind Tb cause remarkably distinct clinical effects. For example, the three MT destabilizing drugs in this study – vinblastine (VB), combretastatin A4 (CA4) and plinabulin (PL) — all depolymerize MTs, and two (CA4 and PL) both bind to the colchicine site. Despite these molecular similarities, they cause markedly different effects in the human body **(Table 1)**. Clinical differences between MT drugs could have several causes: distinct biochemical actions on MTs that trigger unique downstream molecular pathways, heterogeneity in cell-type sensitivities due to differential expression of drug transporters or Tb isotypes (9), differences in pharmacokinetics (PK), or off-target activities. To address the first possibility, different molecular actions, we systematically compared gene expression and signaling responses to three MT destabilizers.

**Table 1.** Clinical comparison of microtubule depolymerizing drugs combretastatin A4, vinblastine, and plinabulin. This table compares three diverse microtubule destabilizing drugs (CA4, VB, and PL) according to key clinical characteristics. Tubulin binding site refers to the known site that the drug binds in either tubulin or the microtubule lattice. Drug class is defined by how the drug is classed in the clinic or in clinical trials. FDA approval indicates whether the U.S. Food & Drug Administration has approved the drug for clinical use. Primary therapeutic use and primary target tissues are brief summaries of drug labels, off-label use, clinical trial aims, and discussions with clinicians. VDA: Vascular disrupting agent; MT: microtubule; FDA: U.S. Food & Drug Administration.

|  | **Combretastatin A4 (CA4)** | **Vinblastine (VB)** | **Plinabulin (PL)** |
| --- | --- | --- | --- |
| **Tubulin binding site** | Colchicine | Vinca | Colchicine |
| **Drug class** | Anti-tumor VDA (first in class) (47) | Vinca alkaloid (48) | VDA, Selective immunomodulating MT agent (first in class) (49) |
| **FDA approval** | No | Yes | No |
| **Primary therapeutic use** | Selective targeting of tumor endothelial cells (50) | Anti-mitotic effects on tumor cells | Reduction of chemotherapy-induced neutropenia |
| **Primary target tissue** | Solid tumors (endothelium) | Liquid tumors (cancer cells) | Bone marrow (neutrophil progenitors) |

VB is a representative vinca alkaloid, a class that is thought to exert cytotoxic effects on cancer cells (3). CA4, under development as the phosphate ester pro-drug fosbretabulin, is being investigated as a complement to chemotherapy which selectively targets the tumor vasculature rather than cancer cells (10). PL exhibited clinical activity that was unprecedented for a MT drug, boosting white blood cell production and protecting the bone marrow during chemotherapy (11). This unexplained clinical activity of PL is remarkable given that most MT drugs suppress white blood cell production via anti-mitotic effects in the bone marrow (12). Colchicine, the most clinically divergent MT destabilizer, is used to treat inflammation but not cancer (2). We omitted it from our panel due to evidence that its unique profile is caused by selective hepatocyte distribution, a PK effect that is hard to model in culture (13). As comparators, we included the MT stabilizing drug docetaxel (DTXL) and the pro-inflammatory cytokine TNF-α to test for pathway overlap with MT drugs.

We profiled molecular pathway responses to MT drugs in primary human pulmonary microvascular endothelial cells. We chose this cell type because endothelial cells use their MTs to regulate barrier function between the bloodstream and surrounding tissues (14), making them sensitive to MT disruption, and because endothelial cells are highly exposed to drugs which are administered intravenously, as is the case for MT drugs used in cancer. Primary cells often show stronger signaling responses to perturbation than established cell lines, and the slower division rate of primary endothelial cells reduces anti-mitotic actions, which we wanted to avoid in this study. VB, CA4, and PL are all known to impact endothelial cells. VB causes endothelial damage (15), and both CA4 and PL are thought to selectively target tumor endothelium (10, 16).

To broadly characterize molecular pathway changes downstream of MT perturbation, we used mRNA-sequencing (RNA-seq) and phosphoproteomics. Several molecular pathways are already known to respond to MT drugs. In interphase, MT destabilization releases and activates GEFH1 (ARHGEF2) (17, 18), which in turn activates Rho GTPases, ROCK, MAP3K1, ERK and JNK (19, 20). In mitosis, MT destabilization promotes mitotic arrest, leading to mitochondrial damage and apoptosis (21). We collected data at short time points to minimize mitotic perturbation. We found that all three MT destabilizing drugs induced equivalent molecular pathway responses, suggesting that differences in their clinical effects have other causes.

## Results

### Microtubule depolymerization assay

Primary human pulmonary microvascular endothelial cells (HPMECs) from three donors were used after fewer than 9 passages **(Supplementary Table 1)**. Three MT destabilizing drugs (VB, CA4 and PL) and the MT stabilizing drug docetaxel (DTXL) were used at used at 100 nM, chosen to achieve saturation of binding sites based on IC_50_ values from prior experiments and literature **(Supplementary Figure 1)**. TNF-α was used at 1 ng/ml, a concentration known to trigger moderate pro-inflammatory signaling. Tubulin staining was used to confirm the expected effect of the drugs. All three destabilizers caused almost complete MT destabilization by 30 min while DTXL reorganized and bundles MTs. TNF-α caused an increase in MT intensity that we did not pursue further.

### MT destabilizers elicit equivalent transcriptional changes in HPMECs

To assess transcriptional responses, HPMECs were treated with the perturbation panel for 6 hours followed by standard bulk mRNA sequencing analysis (RNA-seq). We first compared gene expression across 6 conditions and 3 donors. Donor variability accounted for 57.7% of the variance, primarily captured by principal components 1 and 2 (PC1/PC2) **(Figure 1A)**. Treatment effects were reflected in principal components 3 and 4 (PC3/PC4). We then performed differential gene expression analysis, comparing each drug to DMSO or drugs to each other. Each destabilizing drug induced substantial transcriptional changes, with over 650 differentially expressed genes (DEGs) compared to DMSO ( | log_2_FC | ≥ 1, p-adjusted < 0.05) **(Figure 1B)**. Strikingly, when the three MT destabilizers were compared to each other, there were no DEGs, indicating that the transcriptional program induced by the three destabilizers was remarkably similar. DTXL causes some regulation of transcription, but much less than the destabilizers. TNF-α also caused large changes, as expected, and there was significant overlap in the DEGs between TNF-α and the destabilizers.

**Figure 1.**
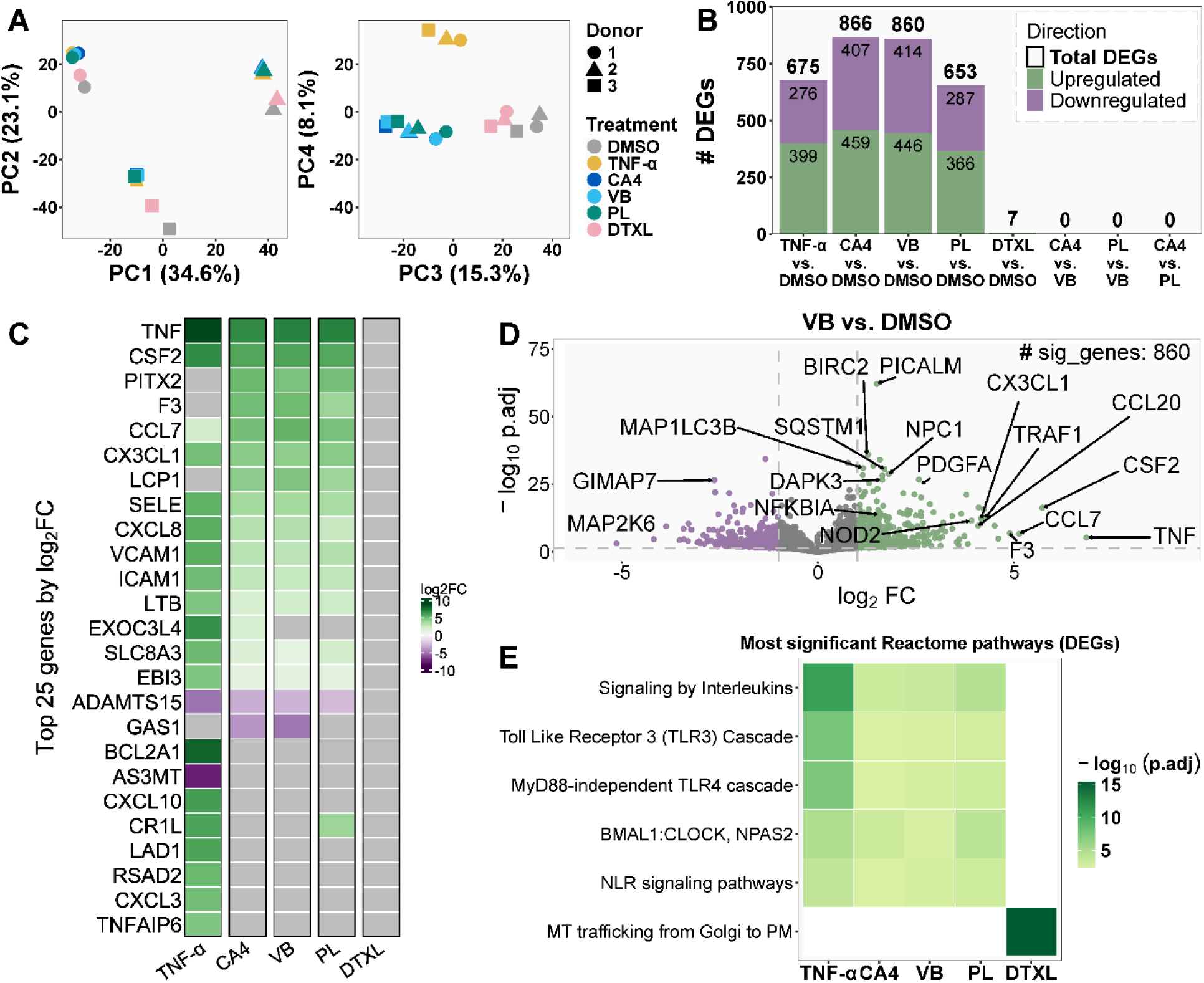
Microtubule destabilizers elicit similar transcriptional responses in HPMECs. Cells were treated for 6 hours with each condition. (A) Principal component analysis of RNA-seq data from HPMECs treated with MT destabilizers. Donors are shown as shapes, treatments as colors. MT destabilizers are colored in shades of blue/green. Percentages indicate proportion of variance explained by each component. (B) Number of differentially expressed genes (DEGs) in HPMECs. Conditions were compared to vehicle (0.1% DMSO), and MT depolymerizers were compared to each other. Genes were classified as differentially expressed if | log_2_FC | > 1 and p-adjusted < 0.05. Directionality reflects changes in the first comparator relative to the second. (C) Top 25 DEGs across all conditions ranked by absolute log_2_FC. DEGs were selected based on the highest absolute log2FC across conditions and displayed in descending order of log2FC for the CA4 condition. Genes not differentially expressed in a given condition are shown in gray. (D) Volcano plot of vinblastine-treated HPMECs compared to baseline. Genes with the highest adjusted p-values (black) or log2FC (blue) are labeled. (E) Reactome pathway analysis of DEGs. The top 3 pathways (by adjusted p-value) were selected for MT depolymerizers, and the top 1 for TNF-α and DTXL. Darker colors indicate more significant adjusted p-values.

We combined functional pathway analysis with manual inspection to analyze the biological implication of the DEGs triggered by all three MT destabilizers in all three donors. We noted prominent upregulation of genes encoding secreted growth factors, cytokines, and adhesion molecules, as well as factors that regulate immunity (NOD2, TRAF1), autophagy (SQSTM1, NPC1, MAP1LC3B), apoptosis (BIRC2, DAPK3), and endocytosis (PICALM) **(Figure 1C-D)**. Downregulated genes included members of the GIMAP family, a diverse set of GTPases implicated in immunity (22, 23). Pathway analysis via Reactome indicate that transcriptional response to MT destabilization significantly aligns with inflammatory signaling pathways **(Figure 1E)**. These results align with prior literature connecting MT depolymerization and inflammatory signaling (24), though we are unable to replicate experiments showing relA/p65 (NF-κB) translocation into the nucleus upon MT depolymerization (25) in our cell models (data not shown). The response to destabilizing drugs was not a pure inflammatory signature, since it differed significantly from TNF-α and included multiple growth factors. We interpret the response as a tissue injury response signature which upregulates inflammatory, wound healing, and pro-thrombotic genes.

To test the generality of the gene expression response to MT destabilizing drugs, we treated quiescent RPE1 cells (immortalized retinal pigment epithelial) with CA4. We noted significant overlaps with the HPMEC response, including shared induction of two potent growth factors, HBEGF and PDGF **(Supplementary Figure 2D)**. Transient MT destabilization was shown to induce cell cycle re-entry in quiescent fibroblasts (26). Induction of growth factors might account for that classic and previously unexplained observation.

### Gene expression response to MT destabilization depends on Rho GTPase activation

To test dependence of the gene expression response on Rho pathway activation, HPMECs were pre-treated with either a pan-Rho inhibitor (a cell permeable from of C3 transferase, RHOi), a ROCK inhibitor (Y-27632, ROCKi), or vehicle, before treatment with 0.1% DMSO or 100 nM VB as a representative destabilizer for 6 hours **(Figure 2A)**. RNA-seq analysis showed that RHOi almost completely abolished the transcriptional response to VB, reducing the number of differentially expressed genes (DEGs) by 98.7% **(Figure 2B)**. In contrast, ROCK inhibition led to a more moderate 27.7% decrease in DEGs. These findings establish Rho GTPase activation as the primary driver of transcriptional changes following MT destabilization. RHOi inhibits RhoA, RhoB, and to a lesser extent RhoC (27). RhoA is the family member most often implicated in response to MT destabilization (18), but RhoB plays important roles in barrier regulation in endothelial cells (28). In contrast, activation of ROCK contributes only partially to the transcriptional signal. These findings are summarized in Figure 3C, where inhibiting Rho GTPases blocks all signaling from MT loss to gene expression, while inhibiting ROCK only has a partial effect.

**Figure 2.**
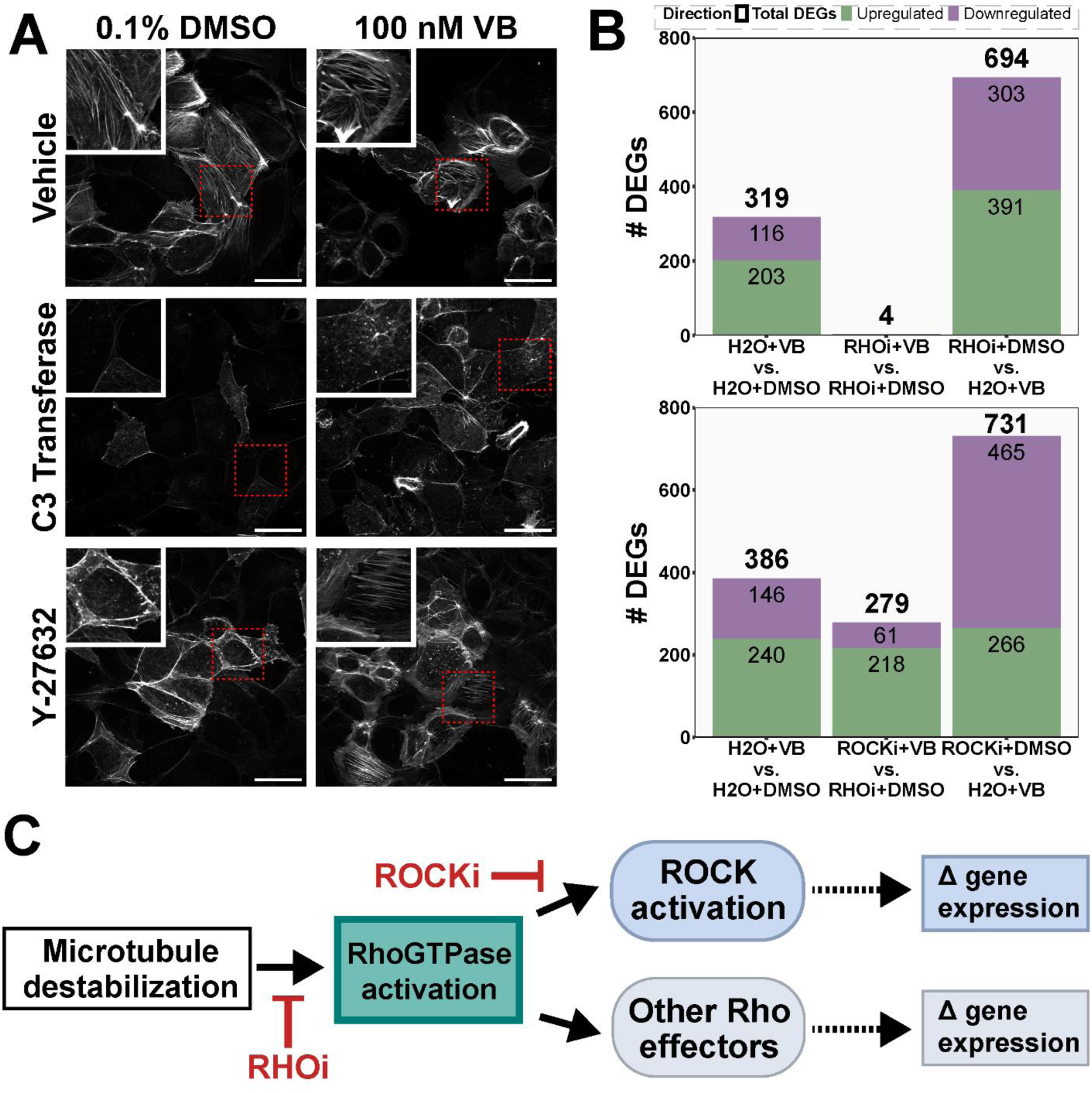
Transcriptional changes are downstream of Rho activation. (A) Change in VB-activated actin stress fibers after Rho and ROCK inhibition. HPMECs were pre-treated with vehicle (H_2_O), 1 μg/ml C3 transferase (Rho inhibitor, RHOi) for 4 hours, or 10 μM Y-27632 (ROCK inhibitor, ROCKi) for 1 hour. After pre-treatment, cells were treated with 0.1% DMSO or 100 nM VB for 30 min. Actin stress fibers were stained with fluorescent phalloidin and imaged on a confocal microscope. Representative images were selected and scaled by the brightest image. Insets show selected regions (red boxes) at higher magnification. Scale bar = 50 μm. (B) Changes in DEGs after Rho and ROCK inhibition. Following pre-treatment, cells were treated with 0.1% DMSO or 100 nM VB for 6 hours, with a booster of 0.5 μg/ml C3 transferase for RHOi. DEGs are genes with | log_2_FC | > 1 and p-adjusted < 0.05. Comparison between perturbation+DMSO and H2O+VB indicates changes in gene expression are not due to the perturbation having a similar effect as MT depolymerization. (C) Model for MT loss-dependent changes in gene expression. MT destabilization activates Rho GTPases, which signal partially through ROCK as well as other effectors. ROCKi leads to a reduction in ROCK-dependent downstream gene expression, while RHOi inhibits changes through all arms of the pathway, completely abolishing changes in gene expression induced by MT destabilization.

**Figure 3.**
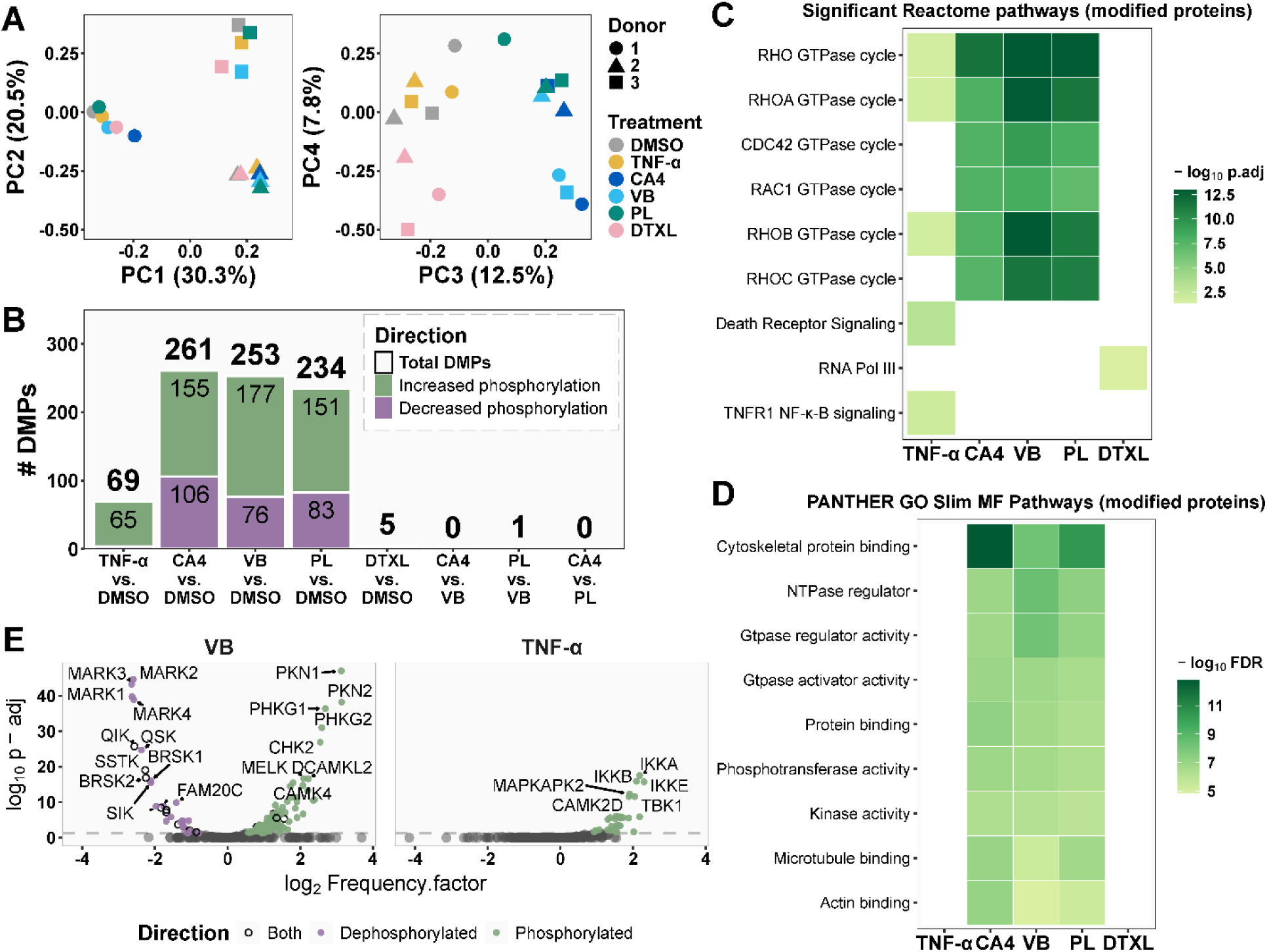
Differentially modified phosphosites (DMPs) are similar across MT destabilizers. (A) Principal components of phosphoproteomic data for short-term MT destabilization in HPMECs. Donors are represented by different shapes, and treatments correspond to different colors. (B) Number of DMPs after 30 min of treatment. Each condition was compared to vehicle (0.1% DMSO), and MT destabilizers were compared to each other. Modified peptides were classified as significantly differentially phosphorylated if | log_2_FC | > 1 and p-adjusted < 0.05. (C-D) Functional pathway analysis of differentially modified proteins. After DMPs were identified, the names of modified proteins were analyzed using Reactome and PANTHER GO Slim. For MT depolymerizers, the top 5 pathways from Reactome and the top 7 pathways from PANTHER were selected, based on adjusted p-value. For TNF-α, the top 2 pathways were included, while for DTXL, only the top pathway was included. (E) Predicted kinase activity in HPMECs treated with VB or TNF-α for 30 min. Centered phosphosites were submitted to the Kinase Library’s updated library to predict kinase activity based on substrate modifications. VB was selected as a representative depolymerizer. Kinases with phosphorylated substrates are shown in green, dephosphorylated substrates in purple, and those with both phosphorylated and dephosphorylated modifications in an open black circle.

### Microtubule destabilizing drugs elicit equivalent phosphorylation response

We next investigated whether the clinical differences may arise from distinct signaling events more proximal to MTs, even if these ultimately converge on a common transcriptional response. HPMECs were treated with the MT perturbation panel for 30 minutes to achieve near-complete MT destabilization, and phosphorylation changes were assessed using quantitative phosphoproteomics. This short time point was chosen to capture immediate signaling effects before gene expression changed cell physiology.

Donor variability accounted for 50.8% of the total variance in the phosphoproteomic data, while treatment effects specifically distinguishing MT destabilizers from other conditions explained the next 12.5% in the 3rd principal component **(Figure 3A)**. Analysis of changes that were statistically significant across three donors showed that MT destabilizers caused differential modification of ∼230 phosphosites, compared to ∼70 with TNF-α and only 5 with DTXL **(Figure 3B)**. No significant differences were detected when comparing the three MT destabilizers to each other. Although the gene expression program triggered by MT destabilizers and TNF-α triggered overlapped significantly, there was no overlap in phosphosite modifications. This shows that the signaling pathways differ and only converge at the level of transcription. Interestingly, the Ser1459 phosphosite on SOGA1/MTCL2 showed opposite regulation, being dephosphorylated by MT destabilizers and phosphorylated by the MT stabilizer DTXL. Treating the three donors as experimental replicates for scoring significance was a particularly stringent cut-off for phosphosite changes. We suspect that additional phosphosite changes in the primary data may be biologically meaningful and refer interested readers to the primary data.

We used Reactome and PANTHER GO Slim to link the proteins being phosphorylated or dephosphorylated upon MT destabilization to the pathways in which they are active. After treatment with all three MT destabilizers, proteins involved in Rho GTPase cycles and cytoskeletal regulation were the most significant categories undergoing modification, including microtubule-associated proteins **(Figure 3C-D, Supplementary Figure 3B)**. This imputation of Rho GTPase regulation at the level of phosphosites is consistent with our observation that all transcriptional regulation depends on Rho GTPases. GEF-H1 (ARHGEF2), which is known to orchestrate signaling responses to MT depolymerization (18, 24), exhibited multiple changes in phosphorylation after MT depolymerization. Some correspond to known regulatory sites in the literature while other sites are novel.

### Imputation of MARK kinase sites in the MT destabilization response

To infer the kinases whose substrate sites were gained or lost following MT destabilization, we mapped differentially modified phosphosites to predicted kinases using the Kinase Library enrichment tool, which bases predictions on kinase preferences for peptide substrates (29, 30). Notably, the predicted kinases were primarily upstream regulators, suggesting that our analysis captures an early stage of the phosphorylation response in HPMECs **(Figure S4)**. Across all MT destabilization treatments, substrates of PKN1/2 and several CAMKs increased in phosphorylation, consistent with Rho activation as a driver of these modifications **(Figure 3E)**.

The most significant kinase imputation was that 45% of the phosphosites lost after MT destabilization are predicted substrates of MARK family kinases (MARK1-4). Some of these predicted substrates are known microtubule-associated proteins (MAPs). This suggests that MT destabilization reduces MARK activity or increases phosphatase activity at MARK sites. MARKs are known to phosphorylate MAPs in a reaction that is stimulated by MTs (31), and sites on MAP4 that were reduced following MT destabilization are annotated MARK sites. Our data suggest that MARKS phosphorylate dozens of proteins in a MT-dependent manner and provide a list of candidate MARK substrates with new entries.

### MT destabilizing drugs differ in washout kinetics

One way to explain clinical differences between drugs might be differences in pharmacokinetics. For example, tissue washout kinetics would affect duration of action in a tissue in a pulsatile exposure regime. To start exploring this hypothesis, we measured washout kinetics in cell culture and found that PL exited cells faster than the other destabilizers **(Figure 4)**. This presumably reflects a faster dissociation rate from Tb, which has not been reported, and might explain PL’s relative lack of bone marrow toxicity.

**Figure 4.**
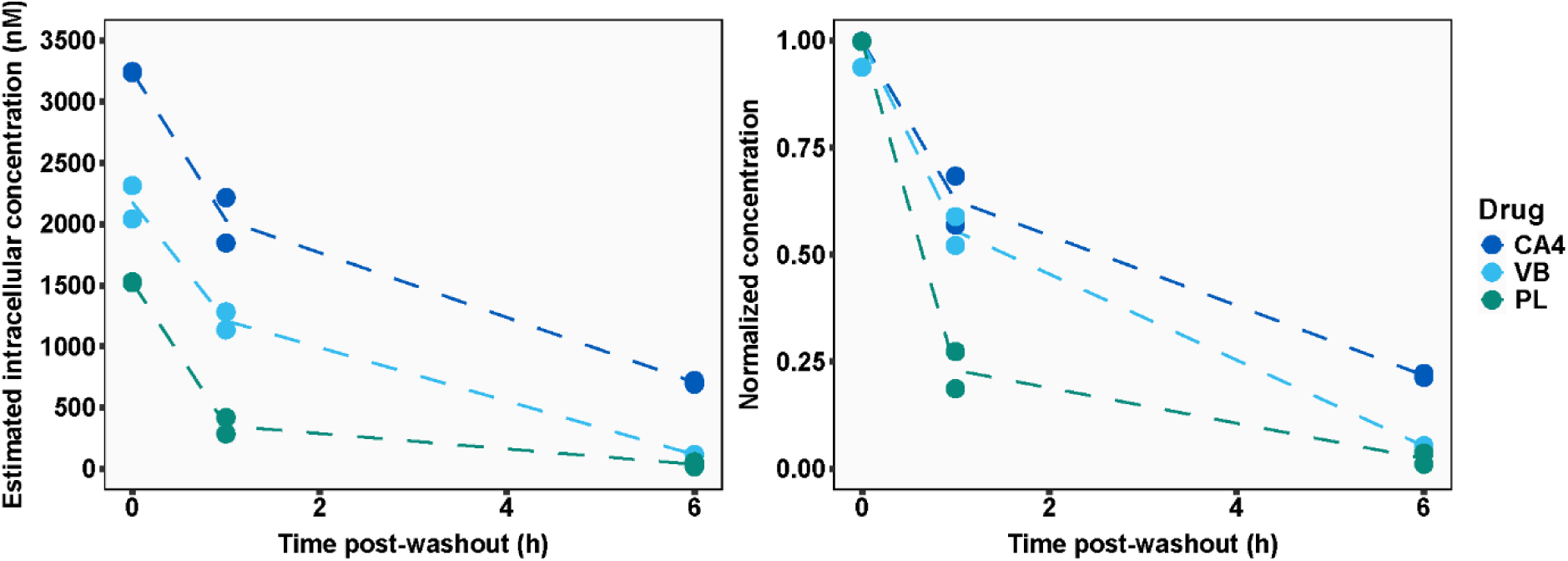
Plinabulin washes out faster from cells than other microtubule destabilizers. HPMECs were incubated with 100 nM of each drug at time 0, followed by four sequential washes at 3, 4, 6, and 9 hours. Cells were harvested at 0, 1, and 6 hours post-initial wash, and intracellular drug concentrations were measured using LC-MS. Drug concentrations are presented as absolute intracellular concentrations (left) and normalized to their initial values (right).

## Discussion

Our primary finding was that three MT destabilizing drugs caused equivalent gene expression and phospho-signaling changes in primary endothelial cells when used at concentrations sufficient to completely depolymerize MTs. This suggests that MT destabilizers, despite differences in biochemical mechanism, converge on shared cellular response programs. This result is expected from a basic cell biology perspective, since all three drugs depolymerize microtubules. It is more surprising from a clinical perspective, particularly for PL, which paradoxically protects the bone marrow.

Previous studies demonstrated that activation of GEF-H1 leads to activation of the Rho family in response to MT destabilization (18). Our work confirms the importance of this pathway by showing that all genes whose expression level changed after 6 hours in VB required Rho activation to respond. GEF-H1 exhibited multiple changes in phosphorylation in our data, more than any other GEF or GAP, which is consistent with it playing a central role. Our work also imputed a major role for regulation of kinase-phosphatase signaling. The strongest imputation was loss of predicted MARK kinase sites when MTs are depolymerized. Based on previous work (31), we hypothesize that the proteins with these sites are phosphorylated by MARKs when both substrate and kinase are MT bound and lost to phosphatase activity when MTs depolymerize. Testing MARK-dependence of these sites will be complicated by co-expression of all four isoforms in HPMECs in our RNA-seq data. Rho GTPases are well known to activate MAPK pathways, but interestingly MAPKs were not strongly imputed by our phosphosite analysis. Instead, the Rho effector kinases PKN1/2 were predicted to be responsible for many of the phosphosites that were added following MT depolymerization. PKN1/2 activation via RhoB has previously been implicated in vascular wall remodeling (32).

If differences in molecular pathway activity do not explain clinical differences between MT destabilizing drugs, what other factors might? PK differences, including tissue washout kinetics, might be one important differential property. Indeed, we found differences in washout rates. MT drugs used in cancer are dosed intermittently, in part to allow bone marrow recovery between doses. Anti-mitotic activity requires that Tb binding sites remain occupied for the duration of cell cycle progression, which is more likely to occur for drugs with slow washout rates. In contrast, the gene expression effects we characterize here are fast and affect all cells in the population at once. We hypothesize that PL induces protective gene expression in the bone marrow following treatment, then washes out quickly which minimizes anti-mitotic toxicity. Another possible difference between drugs that we did not investigate here is cell type tropism. Cell types might differ in expression of drug transporters or tubulin isotypes in a manner that affects their drug sensitivity and different between drugs (13, 33).

Historically, MT drugs have been developed and optimized using largely empirical methods. Our work supports a future where MT drugs can be rationally optimized for different applications based on quantitative understanding of their actions in the human body. Our findings suggest that future drug development efforts should prioritize molecular design elements that influence tissue distribution, cell-type selectivity, and pharmacokinetics, rather than focusing solely on the specifics of drug-Tb interactions, as has traditionally been emphasized.

## Materials and Methods

### Cell culture

All cells were grown in an incubator maintained at 37°C and 5% CO_2_ in a humidified environment. Human pulmonary microvascular endothelial cells (HPMECs, PromoCell #C-12281) from 5 donors were thawed at recommended density of 10,000-20,000 cell/cm^2^. Cells were cultured with complete Endothelial Cell Growth Medium MV (PromoCell #C-22020), passaged with DetachKit (PromoCell #C-41220), and cryopreserved with Freezing Medium Cryo-SFM (PromoCell #C-29910). To properly assess changes in inflammation, we replaced complete media with Endothelial Cell Basal Medium MV (Growth Medium Kit – PromoCell #C-39210) enriched with fetal calf serum (0.02 ml/ml) and human epidermal growth factor (10 ng/ml) for at least 12 hours before all assays to remove heparin and hydrocortisone from the media mixture. HPMECs were only used for experiments up to passage 8. hTert-RPE1 (ATCC, Manassas, VA, USA) cell lines were cultured with a mixture of Dulbecco’s modified medium and F12 (DMEM/F12) supplemented with 10% FBS and 1% penicillin-streptomycin (by volume).

### Drug treatments

Microtubule perturbing agents combretastatin-A4 (CA4, MedChemExpress #HY-N2146), vinblastine sulfate (VB, MedChemExpress #HY-13780), plinabulin (PL, MedChemExpress #HY-14444), and docetaxel (DTXL, MedChemExpress #HY-B0011) were reconstituted as 10 mM stocks in 100% DMSO. TNF-α was dissolved in 0.1% BSA as a 100 µg/ml stock.

Cells were treated with 0.1% DMSO, 100 nM of microtubule drugs, or 1 ng/ml of TNF-α for 6 hours (RNA-seq) or 30 minutes (total proteomics and phospho-proteomics) in media without heparin or hydrocortisone. For Rho or ROCK inhibition, cells were pre-treated with 1 µg/ml of cell-permeable C3 Transferase (Rho Inhibitor I, Cytoskeleton Inc #CT04) for 4 hours, or with 10 µM Y-27632 (ROCK inhibitor, Cayman Chemical #10005583) for 1 hour. Following pre-treatment, cells were treated with 100 nM vinblastine or 0.1% DMSO. An additional 0.5 µg/ml C3 transferase booster dose was added concurrently with vinblastine DMSO for 6 hours to maintain inhibition after Rho re-synthesis. Concentration of Y-27632 was maintained at 10 µM throughout the co-treatment.

### RNA-sequencing and analysis

For HPMECs, a total of 10,000 cells/cm^2^ were seeded into 6 cm dishes for RNA-sequencing or 10 cm dishes for proteomics analysis 48 hours prior to drug treatment until cells were 85% confluent. Media without anti-inflammatories was added to cells 12 hours prior to drug treatment. RPE1s were grown to quiescence by growing them to 100% confluence in 6 well plates, and then incubating them for 5 more days before drug treatment. After drug treatment, cells were washed with 4°C basal media (HPMECs) or PBS (RPE1s) and RNA was isolated with the RNeasy Mini kit (QIAGEN #74104) using the manufacturer’s instructions. Integrity and concentration of isolated RNA were measured with the Agilent 4200 TapeStation, and at least 700 ng of each sample was submitted to AZENTA Life Sciences for library preparation and Illumina sequencing (150 bp paired-end reads, 350 M reads per lane, 20 M reads per sample). A second round of quality control was conducted by AZENTA, and all 60 samples had RNA integrity number (RIN) = 10.0, DV200 ≥ 96.8 (mean = 98.7 across all samples), and a sequencing per sample Mean Quality Score ≥ 38.23 (mean = 38.33 across all samples).

RNA-sequencing analysis was guided by the Harvard Bioinformatics Core Introduction to RNA-sequencing course (34). Adapter reads were trimmed using Trim Galore (v0.6.6) and the quality of the resulting reads was assessed with FastQC (v0.12.1). Expression of RNA transcripts was quantified using Salmon (v1.8.0), and quality of the quantification was reviewed with Qualimap (v2.2.1) after reads were aligned with STAR (2.7.9a). DESeq2 (v1.44.0) (35) was used to obtain normalized count data and to conduct differential gene expression analysis, and package apeglm (v1.26.1) (36) was used to conduct log_2_ shrinkage. Differentially expressed genes were filtered based on effect size (log_2_ fold change > 1) and statistical significance (padj < 0.05). Pathway analysis was performed using Reactome.

### Immunofluorescence staining and microscopy

For imaging experiments, HPMECs were cultured on #1.5 glass coverslips coated with 2% type B gelatin, while RPE1s were cultured on plain glass coverslips. For microtubule staining, cells were fixed with pre-chilled 90% methanol and 50 mM EGTA for 20 minutes at -20°C. For phalloidin staining, cells were first fixed with 4% formaldehyde for 10 minutes, followed by permeabilization in 0.1% Triton X-100 in PBS for 15 minutes at room temperature. After PBS washes, fixed cells were blocked for 1 hour at room temperature using a solution of 2% donkey serum (v/v, Sigma Aldrich #D9663) and 5% BSA (w/v, Sigma Aldrich #A9647). Primary antibodies and dyes, diluted in blocking buffer, were incubated with cells for 1 hour at room temperature at the following concentrations: Rat anti-Tyrosinated tubulin (1:200, YL1/2 – Millipore Sigma #MAB1864); Alexa Fluor 488 Phalloidin (1:20, Cell Signaling Technology #8878). Tubulin-stained samples were incubated for 1 hour with secondary Goat anti-Rat IgG Alexa Fluor-647 (1:400, Invitrogen #A-21247). All samples were stained with 1 µg/ml DAPI (Invitrogen #D1306) for 10 minutes and mounted in 90% glycerol with 20 mM Tris (pH 8.0) and 0.5% N-propyl gallate.

Samples were imaged on a Nikon Ti2 microscope with a Yokogawa CSU-X1 spinning disk confocal head, equipped with a Plan Apo 60x/1.4 NA oil-immersion objective. Fluorophores were excited with the corresponding laser lines (Alexa Fluor 488, LUNF-XL 488 laser line; Alexa Fluor 647, LUNF-XL 640 laser line; DAPI, LUNF-XL 405 laser line), and collected with Di01-T405/488/568/647 dichroic mirror and ET525/50m (AF488), ET705/72m (AF647), or ET455/50m (DAPI) emission filter. Images were acquired with a Hamamatsu ORCA-Fusion BT C15440-20UP Digital sCMOS camera without camera binning for optimal resolution. Multiple stage positions were imaged per sample, and representative images of samples were selected for figure presentation. Image sets used for comparison were processed with identical brightness and contrast adjustments using ImageJ (Fiji) software.

### Dose-response survival assay

Cells were seeded 10,000 cells/well in a total volume of 100 µl in opaque 96-well plates opaque plates one day prior to the assay. Drug treatments were randomly dispensed using the HP D300e Digital Dispenser to minimize edge effects. Following 6 hours or 72 hours of incubation with drug, media was aspirated and replaced with 100 µl of fresh media to ensure accurate volume.

Immediately, 100 µl of CellTiter-Glo Luminescent Cell Viability Assay reagent (Promega #G7570) was added to each well. After 10 minutes of incubation, luminescence was measured using the BioTek Synergy H1 plate reader. Background luminescence was subtracted using media-only wells, and treatments were normalized to the mean luminescence of DMSO control wells. All dose-response experiments using HPMECs were performed with at least three replicates per donor, across all three donors.

### Preparation of cell lysates for proteomics

HPMECs were plated in 10 cm tissue culture dishes at a density of 10,000 cells/cm^2^ and allowed to grow for 48 hours until reaching 90% confluency. After 30-minute treatment with microtubule drug panel in anti-inflammatory free media, cells were washed for 5 seconds with 20 mL of 4°C DMEM (no FBS) and then aspirated. This was repeated three times, and then the cells were immediately lysed by addition of 60°C lysis buffer directly onto the tissue culture plate with vigorous scraping. The proteomics lysis buffer consisted of 6 M Guanidine Hydrochloride (Chem-Impex #29913), 2% Hexadecyltrimethylammonium Bromide (Millipore Sigma BioXtra #H9151), 100 mM Na+EPPS pH 8.0 (Millipore Sigma #E1894), 10 mM TCEP HCl (pH = 7.5) (Oakwood Chemical #M02624), and 1 Roche cOmplete tablet (#50855100) per 12.5 mL of lysis buffer.

### Sample preparation for LC-MS for proteomics

Samples underwent sonication for 10 cycles of 15 seconds at an amplitude of 60 while on ice and were clarified by centrifugation at 20k g for 15 minutes at 4°C (minimal or no pellet was observed). Reduction of disulfide bonds was achieved by adding 5 mM Dithiothreitol (DTT) (1 M stock in water) (Oakwood Chemical #3483-12-3) and incubating at 60°C for 15 minutes. Alkylation of cysteines was performed by adding 50 mM iodoacetamide (BioUltra, Millipore Sigma #I1149) (1 M stock in anhydrous N, N-Dimethylformamide (DMF) (Millipore Sigma #227056)) and incubating in the dark for 1 hour. Alkylation was stopped by adding 30 mM DTT. Protein precipitation was carried out using a chloroform/methanol method as previously described (37). The protein disc was resuspended at a concentration of approximately 2 µg/µL in 6 M GuHCl / 10 mM EPPS pH 8.5. 200 micrograms of protein per sample was diluted to 2 M GuHCl with 10 mM EPPS pH 8.5 and digested for 14 hours using LysC (Wako) at a concentration of 20 ng/µL. Samples were further diluted to 0.5 M GuHCl, followed by the addition of an additional 20 ng/µL LysC and 10 ng/µL Trypsin (sequencing-grade modified, Promega #V5111) before incubating at 37°C for 16 hours. Solvent was evaporated in vacuo, and samples were resuspended in 500 mM EPPS (pH 8.0) at a concentration of ∼1 µg/µL. 200 micrograms of peptides were labeled by adding 40 µL of TMTPro (Thermo #A44520) (20 µg/µL in anhydrous acetonitrile (ACN)) and incubating at room temperature for 2 hours. The reaction was quenched by adding 30 µL of 5% hydroxylamine (Millipore Sigma #438227). The samples were pooled, and solvent removed in vacuo. The combined samples were acidified to pH ≤ 1 with TFA, clarified by centrifugation at 20k g for 10 minutes at 4°C, and desalted using a Sep-Pak (Waters). The Sep-Pak was conditioned with 1 mL methanol, followed by 1 mL of 1% formic acid (FA), after which the sample was added. The Sep-Pak was washed twice with 1 mL of 1% FA and eluted into a 2 mL Eppendorf tube with 1 mL of 35% ACN/1% FA. The solvent was removed in vacuo, and 10% of the sample was subjected to bulk protein level proteomics by resuspending the sample in 140 µL of 10 mM ammonium bicarbonate (pH 8.0) and clarified by ultracentrifugation at 70,000 rpm using a TLA-100 rotor for 30 minutes at 4°C. A total of 100 µL of supernatant was fractionated via reverse-phase chromatography as previously described (38) into a 96-well plate, yielding 24 fractions, 6 of which were desalted using stage-tips (39). Approximately 10 µg of peptides per fraction were then subjected to LC-MS/MS analysis.

Phosphoproteomics was done as above except after the Sep-Pak, the remaining 90% of the sample (∼3.2 mg of labeled and combined peptides) was subjected to a phosphopeptide enrichment according to the manufacturer’s instructions with the Thermo High-Select^TM^ Fe-NTA Phosphopeptide Enrichment Kit (Thermo Fisher Cat # A32992). The solvent form the eluted phosphopeptides was removed in vacuo and the sample was desalted with a stage-tip and subjected to LC-MS/MS analysis.

### LC-MS for proteomics

The analysis was performed on a Thermo Orbitrap Eclipse operated in data-dependent mode, with a survey scan range of 300–1100 m/z at an Orbitrap resolution of 120,000. The maximum injection time was 200 ms with an AGC target of 1e6 and a Lens RF of 60%. The system was connected to a Thermo EASY-nLC 1200 HPLC. Each run lasted 4.5 hours with a gradient ranging from 3% to 18% Solvent B (Solvent A: 0.125% FA, 2% DMSO; Solvent B: 0.125% FA, 80% ACN, 2% DMSO). Chromatography utilized in-house packed 100–360 µm ID/OD microcapillary columns, with ∼0.5–1 cm of Magic C4 resin, followed by 40 cm of 1.9 µm Dr. Maisch GmbH C18-AQ resin (Part #r119.aq.0001) (40). An in-house machined column heater maintained at 60°C was used. Peptides were ionized via a 2.6 kV voltage applied to a PEEK microtee. For bulk protein level proteomics, MS/MS run comprised four separate experiments with varying FAIMS (41) CVs (-32, -42, -52, -62). For phosphoproteomics, 1/6 of the sample was analyzed in 6 different MS runs. A single MS run was done (with all the above parameters) with 4 alternating CVs (-32, -42, -52, -62) and 5 subsequent runs were done with a single, different CV (-35, -40, -45, -50, -55). FAIMS was operated at standard resolution with a carrier gas flow of 3.5 L/min. A dynamic exclusion window of 60 seconds and a filter of +/- 10 ppm was applied. MS/MS isolations were performed using a quadrupole isolation window of 0.4 m/z, and peptides were fragmented with an HCD collision energy of 37%. MS/MS scans were recorded with an Orbitrap resolution of 50,000 and a first mass of 110 m/z. An AGC target of 50,000 was applied, with a maximum injection time of 120 ms. The TMT-Pro reporter ions at low m/z were quantified with a 20 ppm mass tolerance and corrected for isotopic impurities as described (42).

### LC-MS/MS data analysis for proteomics

Data were analyzed and searched using a modified approach as described (42). The Comet search engine replaced Sequest for searches. A precursor peptide mass tolerance of 50 ppm was employed, allowing variable modifications for oxidation of methionine (+15.994914 Da) and deamidation of asparagine (+0.984016 Da). Iodoacetamide modification on cysteine (+57.02146372 Da) and TMT-Pro modification (+304.20660 Da) on n-termini and lysines were set as static modifications. For phosphoproteomics fractions, a variable modification on threonine/serine/tyrosine (+ 79.966331 Da) was allowed. The target-decoy approach was used to control the peptide-level false discovery rate (FDR) at 1% (43, 44), with additional filtering to maintain a 1% FDR at the protein level using protein pickr (45). A minimum Sum Sn of 400 for all conditions was required for protein quantification. TMT-reporter ion data for each protein were normalized to sum to one across conditions. Peptides with an MS1 isolation specificity of ≥ 0.95 were preferentially used for quantitation. Contaminants and decoy matches were excluded.

We analyzed dynamic changes in phosphosites by normalizing our phosphoproteomics data. Specifically, we accounted for changes at the protein level by dividing the relative change observed at each phosphosite by the corresponding change in total protein, when applicable. For downstream analysis, phosphoproteomics data was centered log-ratio transformed prior to principal component analysis and differential modification analysis with limma (v3.60.4). Phosphosites were considered differentially modified with a | log_2_FC | > 1 and an adjusted p-value < 0.05. Reactome and PANTHER GO Slim databases were used for functional pathway analysis.

### Kinase prediction

Centered phosphosites (centered by querying UniProt and manually filling in errors) and the log_2_FC and adjusted p-values associated with them were submitted to authors of The Kinase Library enrichment tool (v1.0.3) (29, 30) with an updated kinase library, now available on PyPi. Phosphosites with adjusted p-values > 0.1 were used as substrates to predict kinase regulators. Enriched phosphosites for each kinase were extracted for analysis of which sites were driving the predictions.

### Washout experiments

HPMECs were seeded in 6-well plates at a density of 10,000–20,000 cells/cm². Once the cells reached approximately 100% confluency, the culture medium was replaced with fresh medium containing 100 nM of CA4, VB, or PL. Cells were incubated with the drugs for 3 hours, followed by four sequential washes with fresh medium at 0, 1, 3, and 6 hours. At 0, 1, and 6 hours post-initial wash, one plate of cells was harvested by adding 1 mL of methanol directly onto the cells and incubating for 30 minutes. The methanol extracts were then collected, dried using a SpeedVac SPD2030 at room temperature under 5.1 torr pressure for 3 hours, and reconstituted in 200 µL of methanol. The prepared samples were analyzed using an Agilent 6530 QTOF mass spectrometer equipped with a 1290 Infinity Binary UPLC system and R18 column. Each sample run was conducted over 10 minutes, employing a gradient elution from 2% to 95% solvent B (Solvent A: 1.0% FA in water; Solvent B: 1.0% FA in ACN). Target compounds CA4, VB, and PL were detected at their respective m/z values: 317.1389, 811.4282, and 337.1664. Chromatographic peaks were integrated to calculate the area under the curve (AUC), which was used to determine the level of each drug in the sample. Drug concentrations were quantified by fitting the LC/MS data to the standard curve generated for each drug. Intracellular drug concentrations were calculated using the estimated total cell volume in each well. The total cell volume was determined by multiplying the bottom surface area of the well (9.6 cm²) by the height of the cell layer (3.7 µm), which was measured using a microscope with z-stack imaging.

### Data and code availability

The RNA-sequencing data sets are publicly available on the GEO database under the GSEA identifiers as follows: GSE286439 (Figure 1), GSE286441 (Supplementary Figure 2), GSE286442 (Figure 2, RHOi), GSE286443 (Figure 2, ROCKi). The mass spectrometry proteomics data have been deposited to the ProteomeXchange Consortium via the PRIDE (46) partner repository with the dataset identifier PXD059740. The source code for analysis can be found on GitHub at http://github.com/lillianhorin/mtd upon publication.

## Acknowledgments and Funding Sources

The authors thank Shannan Ho Sui, Meeta Mistry, and Emma Berdan of the Harvard Chan Bioinformatics Core for assistance with experimental design and RNA-sequencing analysis workflows; the Core for Imaging Technology & Education at Harvard Medical School for assistance with light microscopy; the instructors of the Quantitative Imaging: From Acquisition to Analysis course for microscopy instruction; Kevin Dervishi for technical support; and Franklin Staback Rodríguez for experimental guidance.

L.J.H. was supported by the Ford Foundation Pre-Doctoral Fellowship, the National Science Foundation Graduate Research Fellowships Program (NSF GRFP), and the Howard Hughes Medical Institute Gilliam Fellowship for Advanced Study. This research was funded by National Institutes of Health NIH-GM122784. Consultation and teaching support from the Harvard Chan Bioinformatics Core was funded by the Harvard Medical School Foundry.

## Supplementary Figures and Tables

**Supplementary Table 1.** Human pulmonary microvascular endothelial cell (HPMEC) donor information. The table summarizes characteristics of the three donor cell lines used as biological replicates in the primary figures. All donors provided informed consent, as documented by the vendor, PromoCell. Donor information was provided by PromoCell representatives and the public Certificates of Analysis for each lot.

| **Donor label** | **Doubling time** | **Age** | **Sex** | **Ethnicity** | **Smoker status** |
| --- | --- | --- | --- | --- | --- |
| 1 | 28.7 hours | 57 | Female | Caucasian | Never smoker |
| 2 | 32.7 hours | 54 | Male | Caucasian | Non-smoker |
| 3 | 33.4 hours | 68 | Female | Caucasian | Smoker |

**Supplementary Figure 1.**
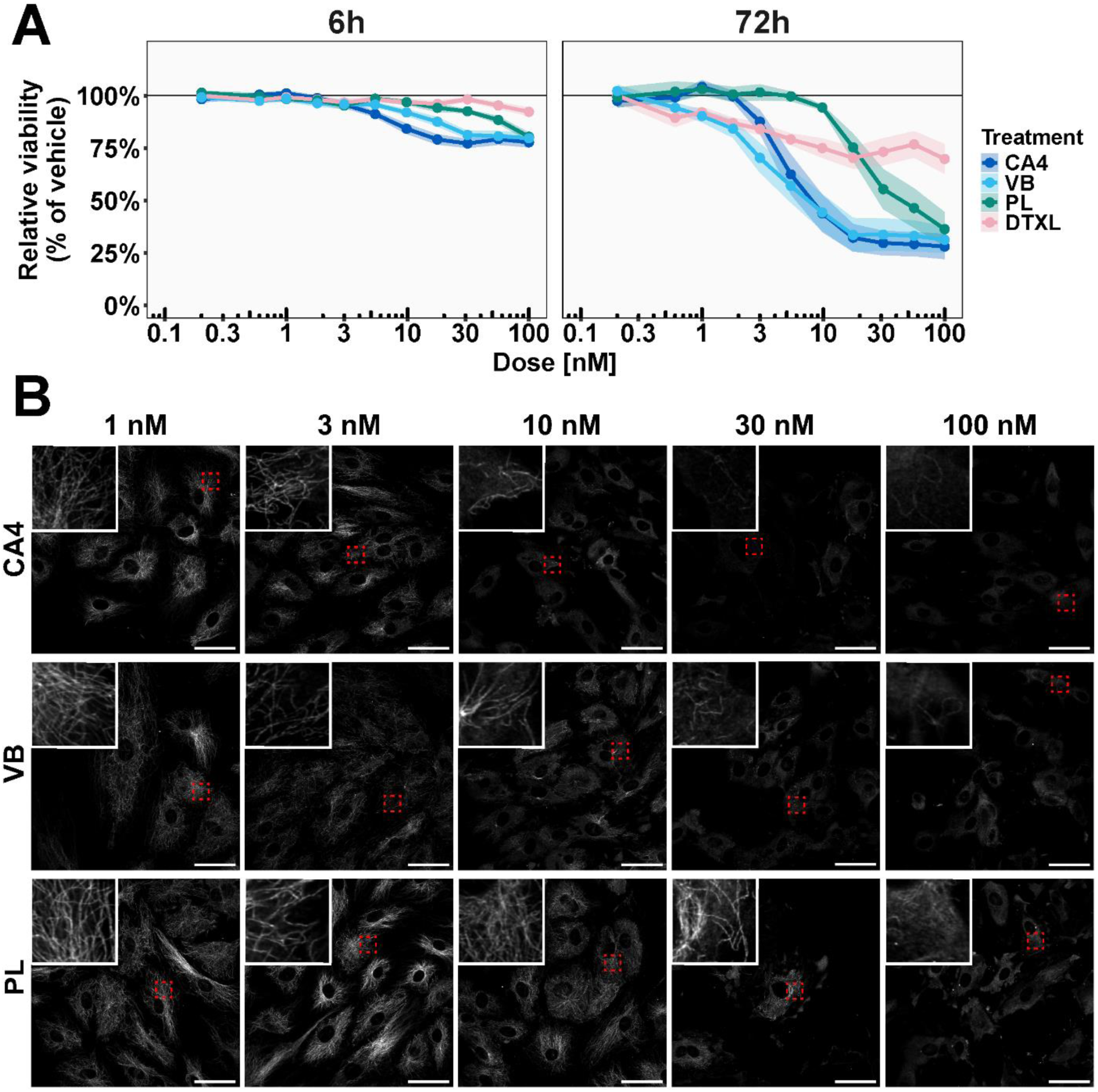
Dose-response of HPMECs treated with MT destabilizers and comparators. (A) Effects of increasing concentrations of microtubule-targeting agents on cell viability. HPMEC viability was assessed after 6 or 72 hours of treatment using the CellTiter-Glo Luminescence assay. Background-subtracted luminescence values were normalized to the 0.1% DMSO control from the respective plate. Error bars represent the standard error of the mean (SEM) from replicates (n = 11). (B) Dose-dependent effects of microtubule destabilizers on tubulin organization in HPMECs. HPMECs were treated with increasing concentrations of MT destabilizers for 1 hour. Tyrosinated tubulin was visualized by immunofluorescence confocal microscopy. Representative images were selected at each concentration and scaled according to the brightest image. Selected regions were highlighted (red boxes) and displayed as insets at higher magnification. Scale bar = 50 μm.

**Supplementary Figure 2.**
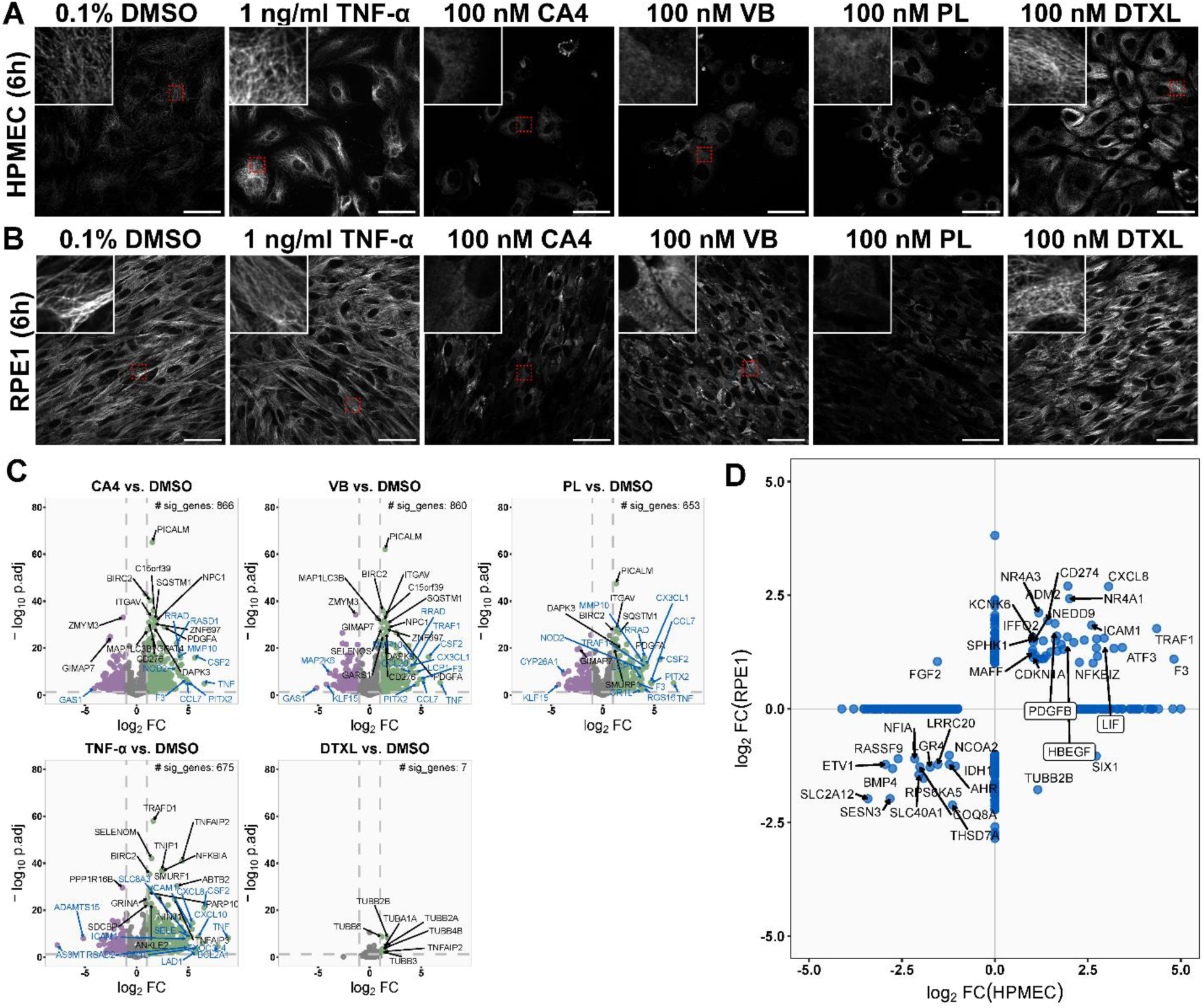
Extended information for RNA-sequencing of HPMECs treated with DMSO, TNF-α, and microtubule-targeting agents for 6 hours. Microtubules were depolymerized after 6 hours in (A) HPMECs and (B) RPE1s. Tyrosinated tubulin was fluorescently stained and imaged with confocal microscopy. Each image is representative of the data and was scaled according to the brightest image. Insets zoom on selected regions (red boxes) at a higher magnification. Scale bars = 50 μm. (C) Volcano plots of DEGs in MT destabilizers and comparators. DEGs with black text are most significant and those with blue text are DEGs with highest log_2_FC. (D) Overlap of differentially expressed genes between HPMEC and RPE1 cells. Overlap was obtained by comparing the Ensembl ID for the genes. Notable growth factors that increase in both cell types are indicated with a black outline.

**Supplementary Figure 3.**
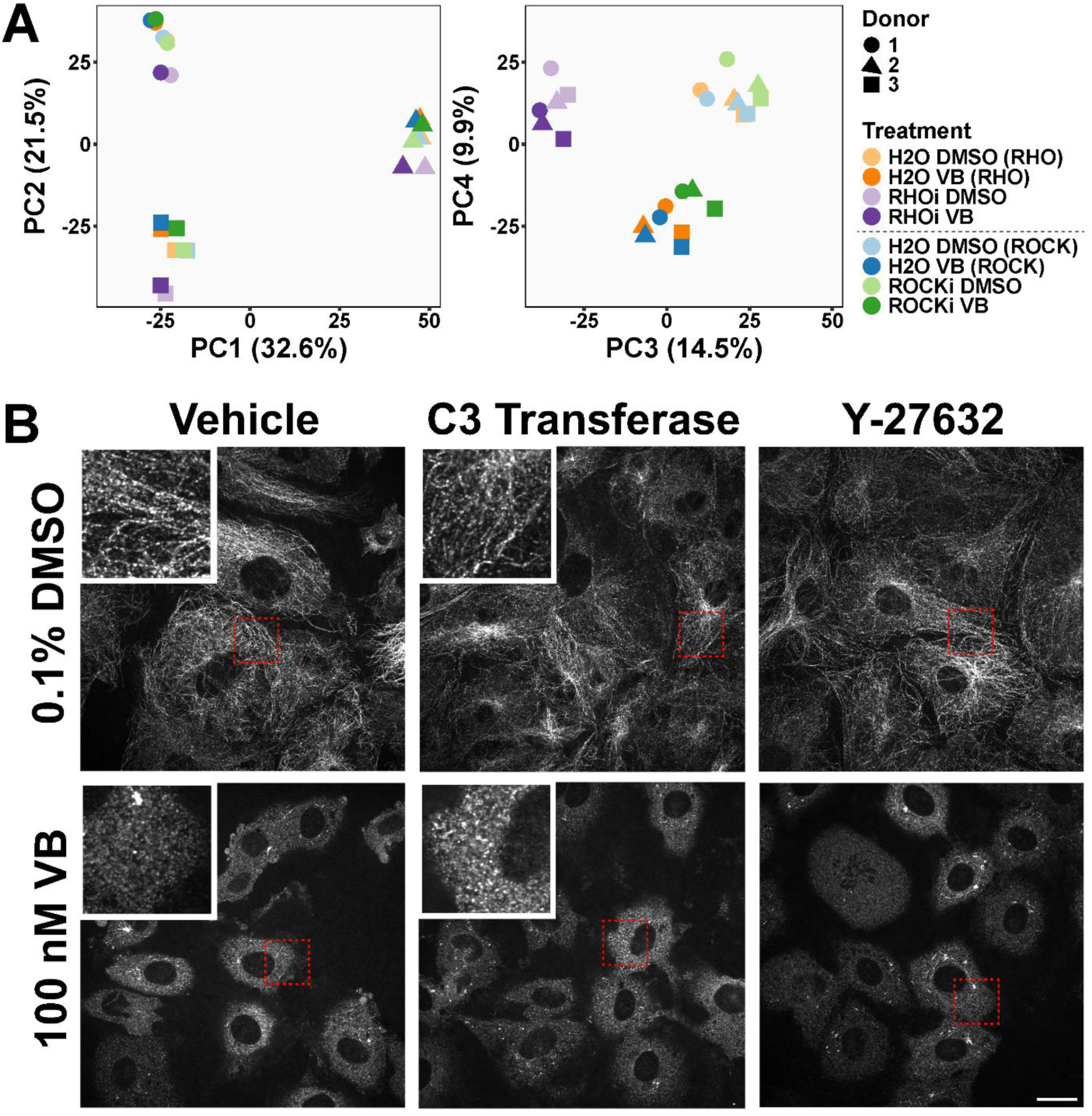
Extended information for RNA-sequencing of Rho/ROCK-inactivated HPMECs. (A) Principle components for samples pre-treated with RHOi/ROCKi and subsequently treated with 0.1% DMSO or 100 nM VB. Shapes correspond to different donors. Different conditions are designated with different colors, and lighter colors correspond to DMSO-treated counterparts. (B) Microtubules after Rho and ROCK inhibition. Cells were treated with 0.1% DMSO or VB for 30 min and imaged by confocal microscopy. Each image was individually scaled for brightness and contrast to enhance visualization and, as such, are not consistent across images. Insets show selected regions (red boxes) at higher magnification. Scale bar = 20 μm.

**Supplementary Figure 4.**
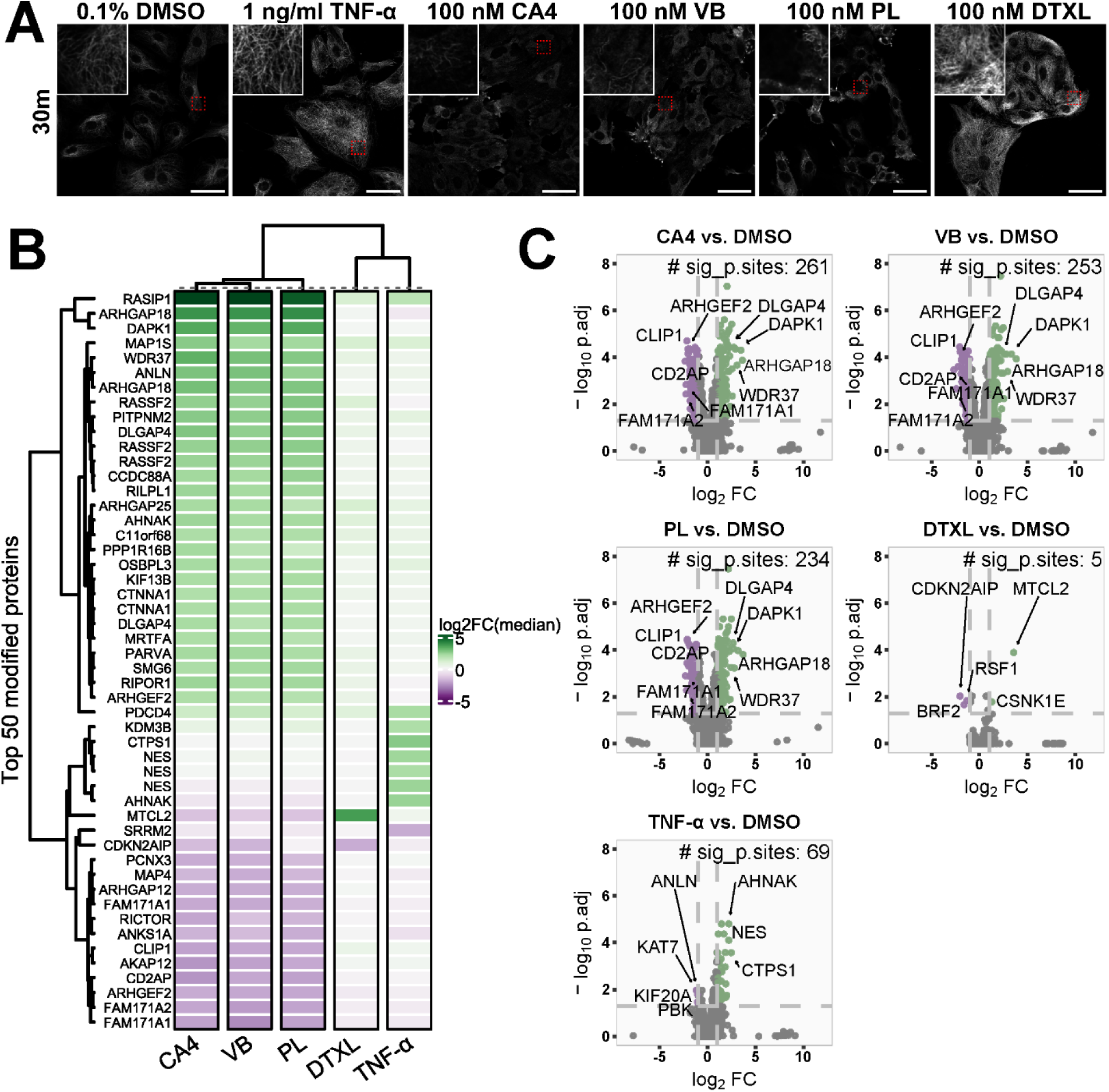
Phosphoproteomics changes in HPMECs after 30 min of MT destabilizer or comparator treatments. (A) Changes in MT polymerization state upon 30 min treatment with drug panel. Each condition is displayed with a representative image and scaled according to the brightest image of the set. Insets are selected regions (red region) at higher magnification. Scale bar = 50 μm. (B) Top 50 most changed modified proteins in panel. The top 50 were selected by rank ordering all phosphosites across the data set by absolute log_2_FC median values and displaying according to log_2_FC of the VB condition, descending. Displayed are the protein names. (C) DMPs in each condition. Significantly dephosphorylated proteins are represented in purple and phosphorylated proteins in green.

**Supplementary Figure 5.**
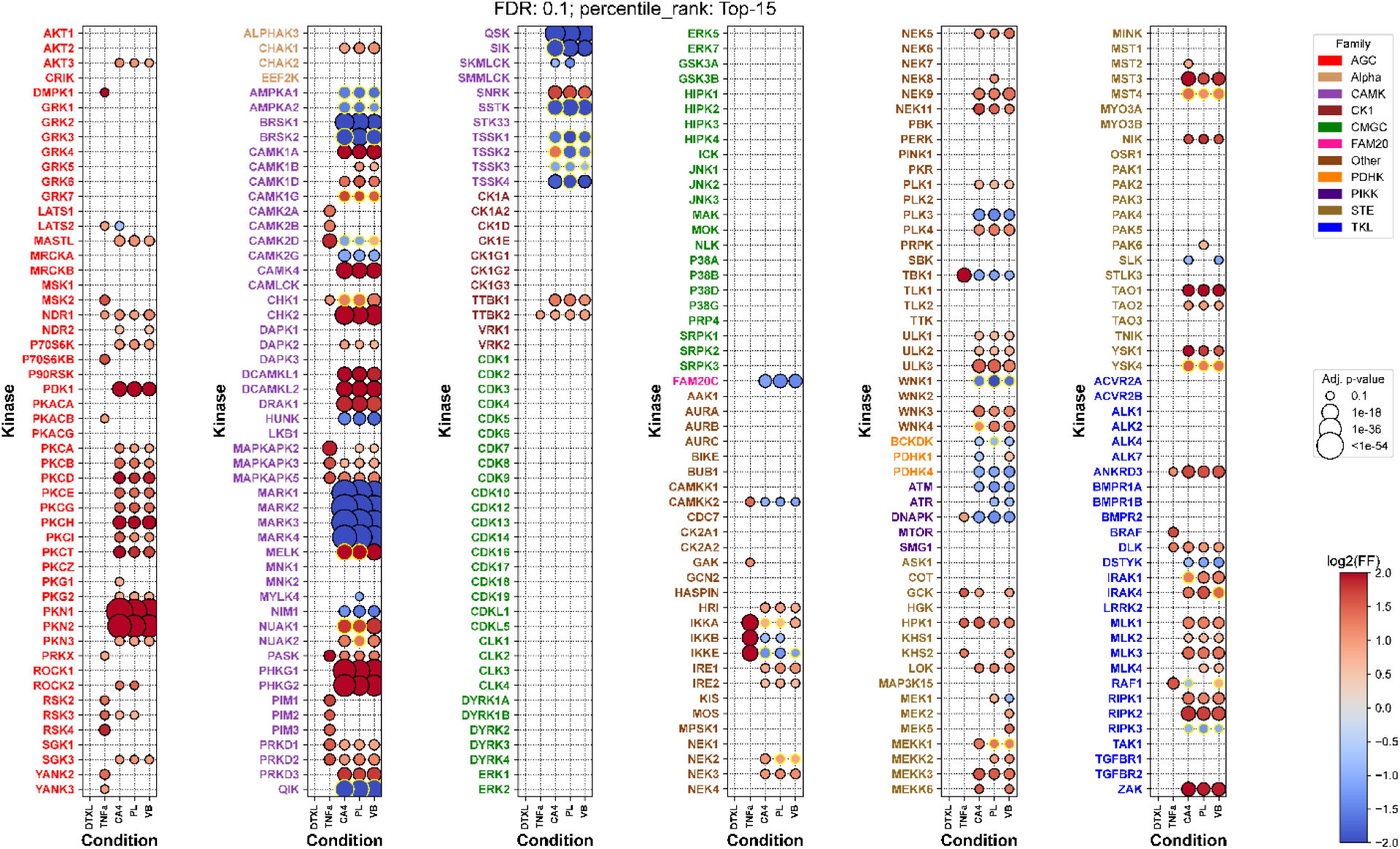
Comprehensive predicted kinases after 30 min of treatment. Each row represents a kinase, grouped by kinase family (color-coded by family type), and each column represents a treatment condition. Circle size corresponds to the adjusted p-value (FDR < 0.1), with larger circles indicating higher significance. The color scale reflects the log₂ frequency factor (FF), indicating motif frequencies. This plot is an output of Kinase Library enrichment tool’s updated package.

